# Integrative, multi-omics, analysis of blood samples improves model predictions: applications to cancer

**DOI:** 10.1101/2020.10.02.299834

**Authors:** Erica Ponzi, Magne Thoresen, Therese Haugdahl Nøst, Kajsa Møllersen

## Abstract

**Background:** Cancer genomic studies often include data collected from several omics platforms. Each omics data source contributes to the understanding of the underlying biological process via source specific (“individual”) patterns of variability. At the same time, statistical associations and potential interactions among the different data sources can reveal signals from common biological processes that might not be identified by single source analyses. These common patterns of variability are referred to as “shared” or “joint”. In this work, we show how the use of joint and individual components can lead to better predictive models, and to a deeper understanding of the biological process at hand. We identify joint and individual contributions of DNA methylation, miRNA and mRNA expression collected from blood samples in a lung cancer case-control study nested within the Norwegian Women and Cancer (NOWAC) cohort study, and we use such components to build prediction models for case-control and metastatic status. To assess the quality of predictions, we compare models based on simultaneous, integrative analysis of multi-source omics data to a standard non-integrative analysis of each single omics dataset, and to penalized regression models. Additionally, we apply the proposed approach to a breast cancer dataset from The Cancer Genome Atlas.

**Results:** Our results show how an integrative analysis that preserves both components of variation is more appropriate than standard multi-omics analyses that are not based on such a distinction. Both joint and individual components are shown to contribute to a better quality of model predictions, and facilitate the interpretation of the underlying biological processes in lung cancer development.

**Conclusion:** In the presence of multiple omics data sources, we recommend the use of data integration techniques that preserve the joint and individual components across the omics sources. We show how the inclusion of such components increases the quality of model predictions of clinical outcomes.

## 1 Background

Cancer studies benefit from the availability of genomic data, also known as omics. The dimensionality of omics data is extremely high, suggesting the application of dimension reduction techniques. Additionally, omics are available across multiple sources (or ‘blocks’) of data, collected on the same organisms or tissues, and measured on different platforms. A comprehensive understanding of the key underlying biological process relies on an integrative approach able to combine the information arising from such multi-source data. To this end, a large number of statistical methods for the simultaneous analysis of multi-omics data have recently been proposed. Multiple reviews of such methods are available, for example in Tseng et al (2015); Huang et al (2017); Rappaport and Ron (2018).

Data integration techniques are often used to identify ‘joint’ (also referred to as ‘common’ or ‘shared’) contributions of the data sources to the observed variation, and their simultaneous effect on the biological process under study. Such patterns of variation arise from the interaction among different omics sources, and may not be detected by a separate analysis of each single source. However, the different data sources do not only contain the joint information, but also independent contributions. The separate analysis of each data source has so far been the most common approach used in the omics context, and knowledge about the individual contributions of each omics source is relevant to the understanding of the biological processes of interest. As a consequence, considering only the joint patterns might also prove insufficient, as it overlooks the heterogeneity among single data sources, and their individual signals from the underlying relevant biological process. An example of this can be seen in genomic studies collecting DNA methylation and gene expression data. It is known that methylation regulates gene expression and that this can cause a non-negligible joint structure across the different data sources. For example, they have been shown to contribute together and jointly relate to the occurrence and characteristics of lung cancer (Heller et al, 2012; Sandanger et al, 2018). On the other hand, methylation and gene expression correlate to these clinical outcomes also through signals that are specific to each omics data source and biologically relevant independently from each other (Yanaihara et al, 2006; Hu and Chen, 2015; Zhang et al, 2016; Baglietto et al, 2017). Therefore, dimension reduction methods that take both joint and individual patterns into account are necessary.

Several methods that have recently been proposed to address this problem are based on matrix factorization. In this framework, each data block is decomposed into three matrices modeling different types of variation, specifically joint variation across the blocks, individual variation for each data block, and residual variation. One such method is JIVE. JIVE stands for Joint and Individual Variance Explained, it was formulated by Lock et al (2013) and, also thanks to its implementation available in R (O’Connell and Lock, 2016), has been used in various medical applications, including clustering of cancer genomics data (Hellton and Thoresen, 2016), multi-source omics data (Kuligowski et al, 2015; Kaplan and Lock, 2017) and imaging and behavioral data (Yu et al, 2017). Although JIVE successfully maintains joint and individual structures, it uses an iterative algorithm and is computationally very intensive. In Feng et al (2018), Angle Based JIVE (aJIVE) was formulated to improve this aspect. It computes the matrix decomposition by using perturbations of the row spaces to identify the joint and individual variation, and results in a much faster implementation than the original JIVE. Besides resulting in a faster implementation of the algorithm, aJIVE provides a more intuitive interpretation of the decomposition, especially in the case of high correlations among individual components (Feng et al, 2018).

Other dimension reduction methods have been extended to the case of multi source data, as for example canonical correlation analysis (CCA) (Hotelling, 1936) or partial least squares (PLS) analysis, which has been further generalized to O2PLS (Trygg and Wold, 2003). A similar method that allows for the presence of multiple data sources is the multiple CCA (Witten and Tibshirani, 2009), but it mainly focuses on the common variation among the components, and seems to neglect the individual contributions of the data sources. An alternative method based on factor analysis has been proposed in Argelaguet et al (2018), and provides a low-dimensional representation of multi-source omics data, although it can fail to detect individual components in the presence of heterogeneous dimensionalities of the single data sources. Other similar approaches to identify both kinds of variation have been proposed, as for example DISCO (Schouteden et al, 2013) and OnPLS (Lofsted et al, 2012). An illustration of these methods and a comparison with JIVE was provided in Måge et al (2019).

Additionally, Principal Component Analysis (PCA) based techniques have been expanded to the case of multi source data. For example, consensus PCA (Westerhuis et al, 1998) consists of PCA on the normalized concatenated data, and distributed PCA (Fan et al, 2019) performs local PCA on the individual data sources and then uses these principal components to estimate a global covariance structure. Integrated PCA (iPCA) is a model based generalization of PCA that decomposes variance into joint and individual variation (Tang and Allen, 2018).

In this work, we focus on prediction models for lung cancer development using both joint and individual components arising from different sources of omics data. We show how the inclusion of both joint and individual components in predictive models leads to a better quality of predictions. The combination of joint and individual components can also facilitate the biological interpretation of the underlying process, although this might still fail as the dimension reduction itself bears the risk of obscuring some relevant information.

We use aJIVE to formulate integrative prediction models in a real data set on lung cancer, identifying individual and joint components across three sources of omics data. We chose to use aJIVE because it inherits a good subspace recovery in comparison to other methods (Måge et al, 2019), as well as the robustness to model misspecification from JIVE, but it also solves the issue of correlated individual subspaces (Feng et al, 2018), and provides a much faster implementation. Furthermore, McCabe et al (2020) show that aJIVE performs best in terms of consistency and lack of overfitting when compared to other integrative methods. We use the aJIVE joint and individual components to build prediction models for lung cancer development.

We evaluate the performance of the proposed models in terms of prediction quality, and we compare them to non-integrative benchmark methods, as well as standard regularized variable selection techniques. Additionally, we show how disentangling joint and individual sources of variation can lead to the identification of biological mechanisms, which would not be highlighted by source-specific analyses.

The data we use stem from a lung cancer case-control study nested within the Norwegian Women and Cancer (NOWAC) cohort study (Lund et al, 2008). The associations among three levels of omics data analyzed in blood samples, specifically DNA methylation, mRNA and miRNA expression, are investigated and their joint and individual contributions are used to predict future cancer cases, and for the characterization of future cancers as metastatic or non-metastatic at diagnosis. We show that both types of components contain information that reveal properties about biological processes and that using joint and individual components results in good model predictions for case-control and metastatic status. We assess the quality of prediction by comparing models based on both joint and individual components to models uniquely based on clinical, patient-level covariates, and most importantly non-integrative models, i. e. based on independent analyses of data from each source.

To further evaluate this approach, we provide an application to a publicly available dataset on breast cancer from The Cancer Genome Atlas (TCGA).

## 2 Methods

### 2.1 Data integration setup

Throughout the manuscript, we will denote each data block with ***X**_κ_*, where *κ* = 1,…,*K* and *K* is the number of data sources used in the study. Each block is a matrix with *n* columns, where *n* is the number of study subjects. The *κ*th matrix ***X**_κ_* has *p_κ_* rows, corresponding to the variables in data source *κ*. The overall dimensionality is denoted as *p* = *p*_1_ +… + *p_K_*. The low-rank decomposition we want to obtain is:

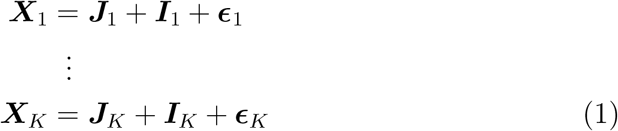

where ***I**_κ_* is the individual component for data block *κ*, ***ϵ**_κ_* is its residual component and

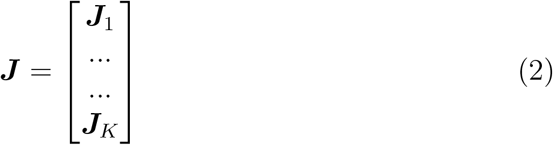

is the joint structure matrix, where each ***J**_κ_* is the submatrix of the joint structure **J** associated with ***X**_κ_*.

### 2.2 Angle Based JIVE

Angle based Joint and Individual Variation Explained (aJIVE) is a variant of the JIVE method, based on perturbation of row subspaces. JIVE aims to minimize the squared residual components *ϵ*_1_,…, *ϵ_K_*, using an iterative algorithm that alternatively estimates the joint and individual components by singular value decomposition (SVD). AJIVE builds on this method but constructs the algorithm in a more efficient and computationally feasible way. The aJIVE algorithm is structured in three phases: First the low-rank approximation of each data block ***X**_κ_* is obtained by SVD. Secondly, the joint structure between the obtained low-rank approximations is extracted by computing the SVD of the stacked row basis matrices. This second phase of the algorithm is based on basic principles of Principal Angle Analysis. Finally, the joint components ***J**_κ_* are obtained by projection of each data block onto the joint basis, while the individual components ***I**_κ_* are calculated by orthonormal basis subtraction.

The first step is based on the choice of the initial ranks for each data block, which are used as a threshold value in the first SVD decomposition of the data blocks. This choice is rather subjective and involves taking into account some bias variance trade-off in the joint signals representation. Although Feng et al (2018) provide guidelines on how to determine the initial ranks, the recommended choice is based on the observation of scree plots, which remains highly subjective. As an alternative, Zhu and Ghodsi (2006) present a choice of initial ranks based on the profile likelihood of the single data blocks.

From the aJIVE decomposition, it is possible to obtain the full matrix representation of the original features, as well as the block specific decompositions of each data source and the common normalized scores. The aJIVE implementation is available in `Matlab` (Jiang, 2018) and `R` (Carmichael, 2019).

### 2.3 Application to the NOWAC data

#### The dataset

The data used in the following analyses stems from blood samples in a lung cancer case-control study nested within the Norwegian Women and Cancer Study (NOWAC) (Lund et al, 2008). All participating subjects are women who did not have a cancer diagnosis at time of blood sampling (2003–2006). The time from blood sampling to cancer diagnosis ranges from 0.3 to 7.9 years, with a median time equal to 4.2 years. The study was designed as a nested case-control study, starting from 125 subjects who developed lung cancer in the NOWAC cohort. One control was randomly chosen for each case from the risk set at the time of cancer diagnosis, following an incidence density sampling scheme. Cases and controls were matched on time since blood sampling and birth year. All participants gave written informed consent and the study was approved by the Regional Committee for Medical and Health Research Ethics and the Norwegian Data Inspectorate. Three levels of omics data are available for *n* = 230 individuals (115 case-control pairs), with numbers of variables respectively equal to *p*_1_ = 485, 512 CpG methylation, *p*_2_ = 11, 610 mRNA expression and *p_3_* = 198 miRNA expression. Information about individual covariates, including age, body mass index (BMI) and smoking habits was also collected for all participants. Outcomes of interest are the classification of case versus control, as well as the characterization of cancers as metastatic or non-metastatic at diagnosis.

#### Filtering and preprocessing

Laboratory processing and microarray analyses for DNA methylation and mRNA expression are described in Sandanger et al (2018). For miRNA, laboratory processing included miRNA isolation and purification from 100 *μl* plasma using the Qiagen miRNeasy Serum/Plasma Kit. Small RNA sequencing libraries were prepared using the NEXTflex small RNA-seq kit v3 (Bioo Scientific, Austin, TX, USA) and sequencing of fragments was performed using a Illumina HiSeq4000 flowcell, according to the manufacturer’s instructions (Illumina, Inc., San Diego, CA, USA), at 50 bp SE, resulting in approximately 7 — 9M reads per sample.

Preprocessing and quality control of methylation data accounted for missing values and intensities below detection thresholds, and included background subtraction and dye bias correction Guida et al (2015). For mRNA data, the probe values were background-corrected and probes reported to have poor quality from Illumina or detected in less than 95% of samples were filtered out Sandanger et al (2018). The filtering of miRNA expressions was based on the counts per million, that is the total read counts of a miRNA divided by the total read counts of the sample and multiplied by 10^6^, and signals having less than one count per million were excluded. Additionally, signals with null reads on more than 5 patients were excluded.

Because of the high computational requirements, we reduced the number of mRNA expressions to *p*_2_ = 5, 000, by selecting the variables with higher variance. We then reduced the number of methylation sites by selecting the CpGs located on the same genes as the filtered mRNAs, as well as the 10,000 CpGs with highest variance. Among these, we excluded CpGs with more than 40% missing data, as well as CpGs with extreme M-values (|M| > 3, see Zhang et al (2012); Ma et al (2014)). This resulted in *p*_1_ = 26, 706. All *p_3_* = 198 available miRNAs were analysed. Other possible filtering criteria have been considered and are described in the discussion of the paper. We used log2 transformed expressions for both mRNA and miRNA, and M-values for methylation (Du et al, 2010). We accounted for missing values in the data by using `SVDmiss`, as suggested in Lock et al (2013). The data was mean-centered. Thanks to the insensitivity of aJIVE to scale heterogeneity, scaling was not performed in the data normalization stage.

#### aJIVE

We performed aJIVE on the three levels of omics data. The initial ranks were selected by maximizing the profile likelihood (Zhu and Ghodsi, 2006), but different choices of initial ranks were also explored and results did not change substantially.

Joint and individual components were used in prediction models. The outcomes of interest were the occurrence of lung cancer (yes/no) and metastasis (yes/no).

We fitted logistic models on each outcome using joint and individual components as explanatory variables, in addition to age, BMI and smoking. These models were compared in terms of AUC with the respective models with only age, BMI and smoking as covariates. To assess the performance of the models, we measured the average AUC in a 10-fold cross-validation. We compared these with a non-integrative analysis, obtained by performing PCA separately on each single data source. We fitted a model on the first principal components (PCs) of each data source, and on the same clinical covariates. We chose to include five PCs for each data source, based on the variance explained by the first PCs and on the analysis of screeplots. We included the same numbers of individual components in the integrative model described above.

To provide a comparison with a standard supervised prediction method, we ran a lasso procedure that selects signals from all three omics layers and used it to predict the two outcomes of interest. We used 10-fold crossvalidation on 2/3 of the data points to select the optimal penalty parameter and we used the fitted lasso model to predict case-control status and metastasis. We ensured the inclusion of the clinical covariates in the lasso models by fixing the corresponding penalty parameters to 0 for age, BMI and smoking status. We compare the quality of model prediction in terms of average AUC across 50 repeats of this procedure.

In addition, we used a random forest of 1, 000 trees to predict case vs control on the basis of joint and individual components, and patient covariates as above (age, BMI, smoking). We extracted the AUCs and the out of the bag (OOB) classification errors from the random forest, and we ranked all the variables in importance on the basis of their mean decrease in gini index (Jiang et al, 2009).

### 2.4 Application to the TCGA data

To assess the predictive performance also in another dataset, we illustrate an application to a subset of data generated by The Cancer Genome Atlas (TCGA Research Network, https://www.cancer.gov/tcga), and used in the mixOmics project Rohart et al (2017).

Records are included for 379 patients, and consist of 2,000 CpGs, 2,000 mRNA and 184 miRNA expression. We use methyaltion, mRNA, and miRNA expression data to explore shared and data-specific components of variation via aJIVE. The joint and individual contributions are used to predict tumor subtypes, specifically a four level classification into Basal, Her2, LumA and LumB breast cancer. The original classification of these subtypes is based on levels of mRNA expression Sørlie et al (2001). We build a prediction model based on joint and individual components, and compare it to a non-integrative model, i. e. based on independent analyses of data from each source.

## 3 Results

### 3.1 Application to the NOWAC data

#### 3.1.1 aJIVE

Using initial ranks obtained with the profile likelihood method resulted in a joint rank equal to 5, and individual ranks respectively equal to 67, 10 and 9. Figure 1 reports the proportions of variance explained that are due to the joint, individual and residual components.

**Figure 1:**
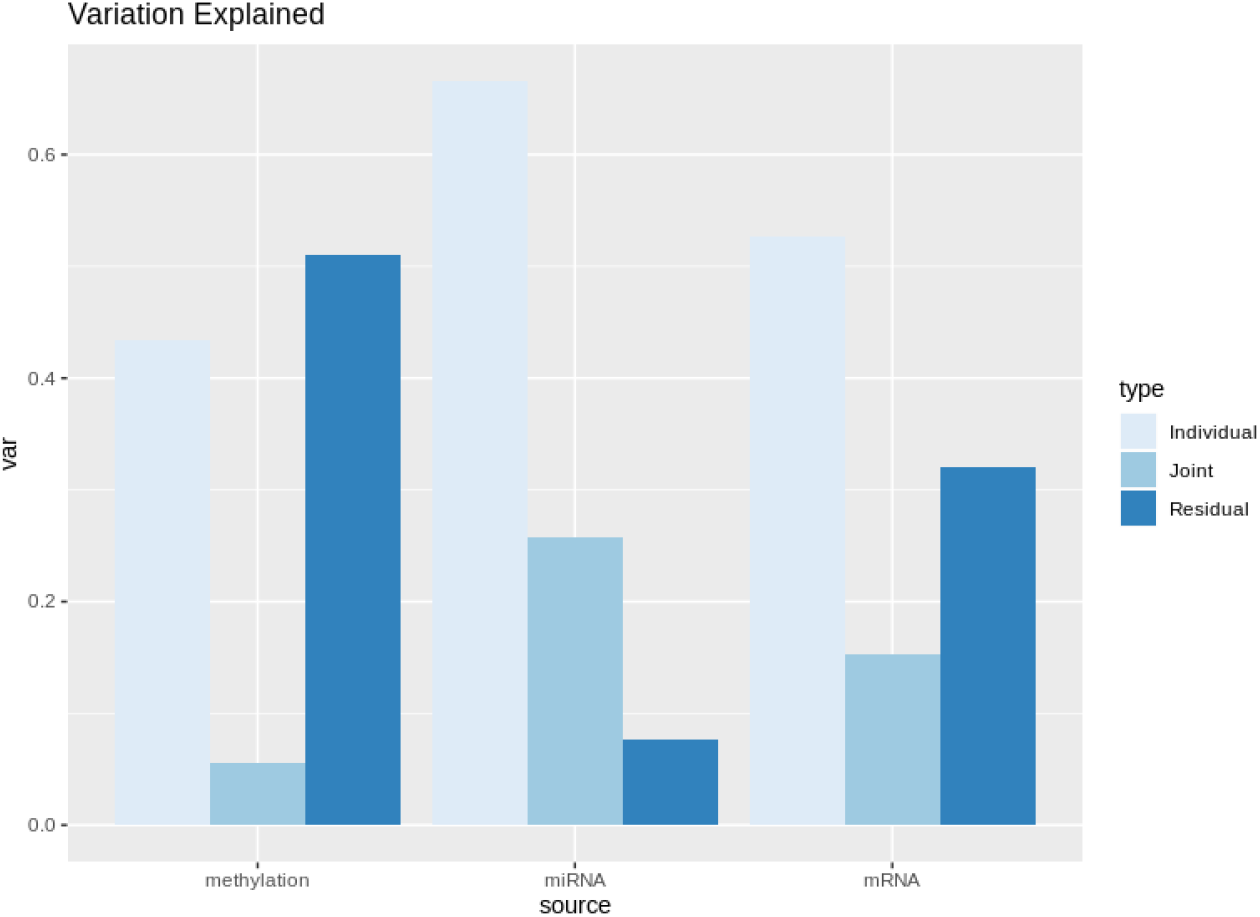
Joint and individual proportions of variance explained in the NOWAC dataset. The individual component is prevalent for all three datasets, especially for methylation. The joint component is relevant for mRNA and miRNA.

Estimated proportions of variance explained with different choices of initial ranks are stable and reported in Supplementary File 2.

#### 3.1.2 Prediction models

Figure 2 reports the in-sample ROC curves relative to the logistic models fitted on the joint and individual components estimated by aJIVE. The model with only patient covariates (age, BMI and smoking) as explanatory variables and the full, integrative model are reported. The integrative model is fitted using patient covariates, aJIVE joint components and first five aJIVE individual components for each data source as explanatory variables. These are compared to non-integrative models, using the first five individual PCs obtained for each dataset separately, in addition to the same covariates. In the prediction of both outcomes, the integrative model shows the highest AUCs, showing how the combination of both kinds of components results in better model predictions. In particular, the integrative models perform better than a non-integrative analysis based on source specific PCs. Additionally, the omics data contribute substantially to the predictions, and result in considerably better prediction quality than patient covariates alone.

**Figure 2:**
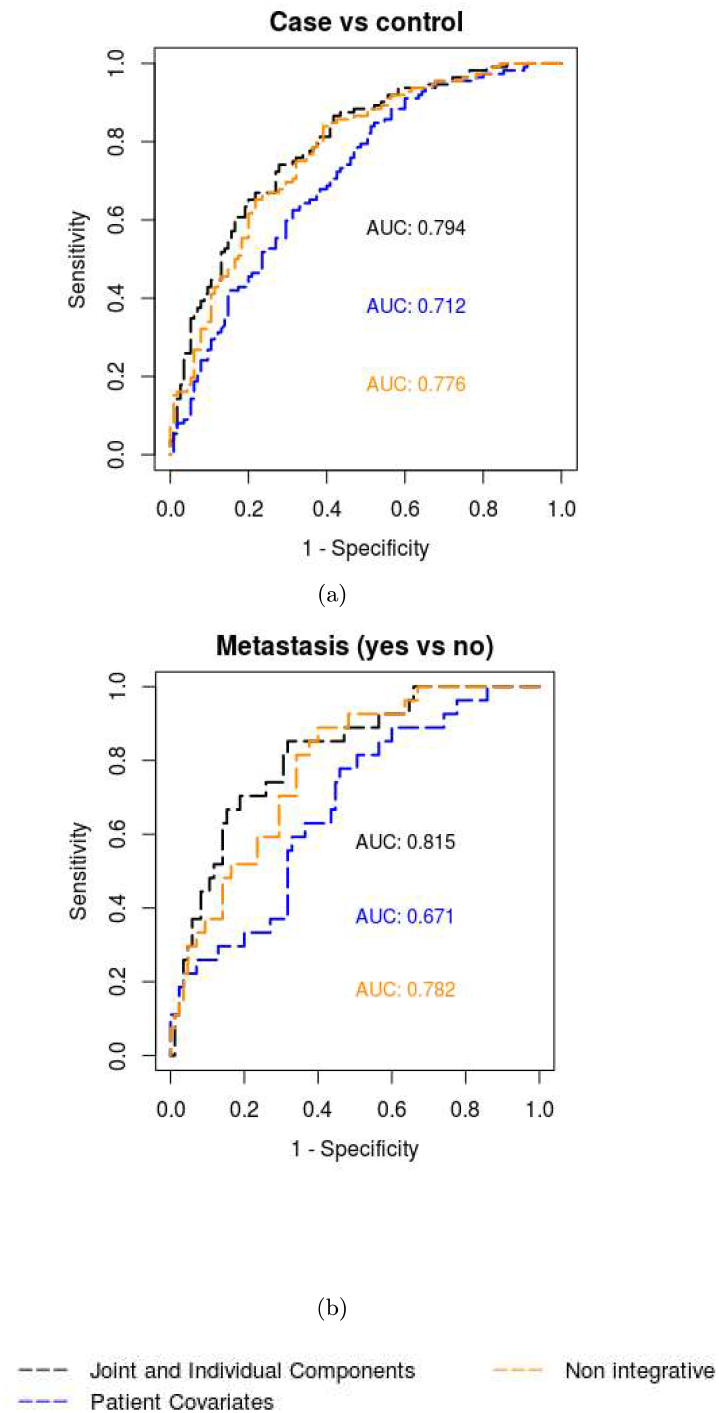
ROC curves from logistic prediction models. a) reports the ROC curves and their AUCs for the prediction models on case vs control, b) reports the ROC curves and their AUCs for the prediction models on metastasis status. The integrative models are fitted on the joint and individual components extracted from aJIVE, while the non-integrative models are fitted on the first principal components obtained separately for each omics source.

A 10-fold cross-validation was used to validate the in-sample results for each outcome, and shows a considerable improvement for the aJIVE integrative model, when compared to non-integrative analysis. In the ROC studies from cross-validation, the integrative models based on the aJIVE components improve the prediction for both case-control and metastasis status. The mean AUCs for the integrative models are 0.69 and 0.70, for case-control and metastasis status respectively. The mean AUC of the non-integrative model, based on the single data source PCAs and the clinical covariates, is respectively 0.65 and 0.63. In the prediction of both outcomes, the aJIVE integrative models perform better than the non-integrative analysis.

For comparison, as mentioned above, we also ran lasso models on the two outcomes. The mean AUCs obtained by the lasso procedure are 0.69 and 0.61 for case-control status and metastastasis, respectively.

Table 1 reports accuracy and OOB classification error for the random forests, as well as the mean AUCs. For case-control status, the aJIVE based model improves the quality of predictions, both in terms of accuracy and AUC, compared to the non-integrative model. The difference from the logistic models with cross-validation can be due to the instability of the random forest, and to the limited sample size. We do not report the random forests results for metastasis because they are highly unstable and the accuracy is very low, most likely due to the even more limited sample size, that is only 125 (only cases) for the metastasis classification.

**Table 1:**
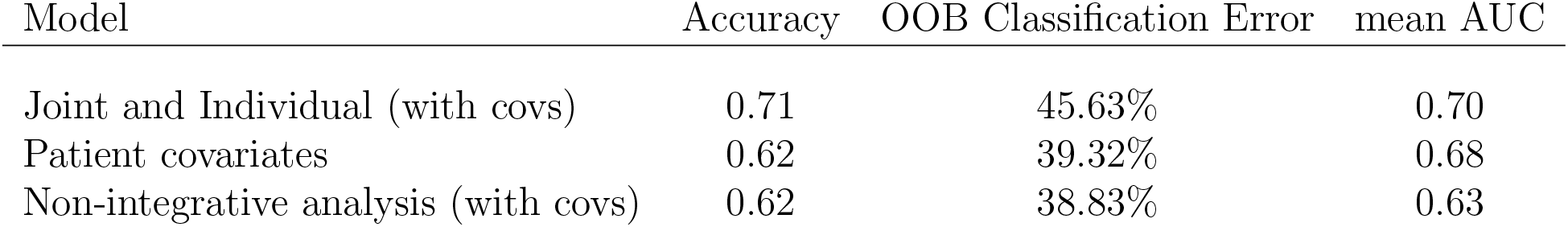
Random forest diagnostics for prediction models of case-control status in the NOWAC dataset.

Figure 3 shows the first ten variables ranked by variable importance in the integrative model for case-control status. One joint component and three individual components appear among the first five variables when ranked for variable importance in the random forest prediction.

**Figure 3:**
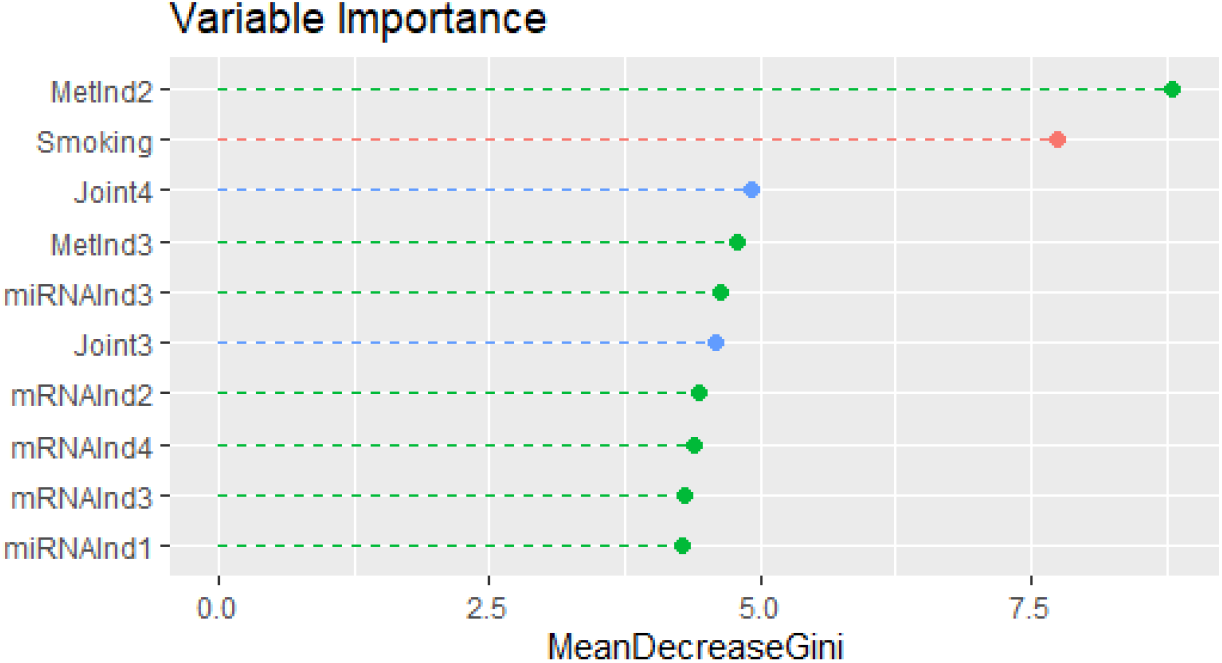
Variable importance plot from random forest on case vs control in the NOWAC dataset. First ten variables ranked by variable importance (in terms of mean Gini index) in the full integrative model for case vs control in the NOWAC dataset. `Joint`_*i*_ denotes the *i*—th joint component estimates by aJIVE, while `MetInd`_*i*_, `mRNAInd`_*i*_; and `miRNAInd`_*i*_ are the *i*—th individual components estimated by aJIVE for methylation, mRNA and miRNA respectively.

#### 3.1.3 Biological interpretation

To investigate the biological processes indicated in the most influential components, we extracted the top genetic features from any omic level that contributed to the first ten variables identified by the random forests. We investigated the omics signals with the highest contribution in terms of loadings estimated by aJIVE, for each component identified by the random forests. Among the mRNAs with the highest contribution, 13 have earlier been identified in conditional logistic regression analyses of metastatic cases sampled within 3 years of their diagnosis as compared to their controls Nøst et al (2020). Additionally, 11 genes overlapped with the top genes identified in the analyses of all case-control pairs independent of metastatic status. We used the Bioconductor package “clusterProfiler” Yu et al (2012) to conduct functional enrichment analyses of GO(BP) categories for these genes, and identified the following ontology categories: inflammatory response, peptide secretion, innate immune response, positive regulation of DNA-binding transcription factor activity, protein secretion, establishment of protein localization to extracellular region. Inflammatory response was identified in Sandanger et al (2018) in the non-smoking related integrative analysis of DNA methylation and gene expression. Among the miRNAs with highest contributions, 80 are significantly associated with the classification of cases vs controls, respectively 36 from the first individual component and 44 from the third individual component. Using the Bioconductor package multiMiR and validated databases therein Ru et al (2014), 55,267 miRNA-gene target interactions were identified for the 36 miRNAs with the highest contribution to the first individual miRNA component. Among the known gene targets for these miRNAs, there were ten (S100A12, MX2, EIF2AK2, TNFSF13B, FFAR2, IL1RN, ANXA3, CCR1, TNFAIP6, TLR5) that were among the mRNA with the highest contributions to the aJIVE mRNA component (“mRNAInd3”). Correspondingly, 32,707 miRNA-target interactions were identified for the 44 miRNAs in the third individual miRNA component. Among these, three (IL1RN, FFAR2, EIF2AK2) were among the mRNA with the highest contribution to the aJIVE mRNA component (“mRNAInd3”).

### 3.2 Application to the TCGA data

#### 3.2.1 aJIVE

Using initial ranks obtained with the profile likelihood method resulted in a joint rank equal to 4, and individual ranks equal to 2, 6 and 11, respectively for methylation, mRNA and miRNA. Figure 4 reports the proportions of variance explained that are due to the joint, individual and residual components. The joint components explain about 30% of the variation in the datasets, and the individual contributions are limited to about 25%, leaving a high contribution to the residual components.

**Figure 4:**
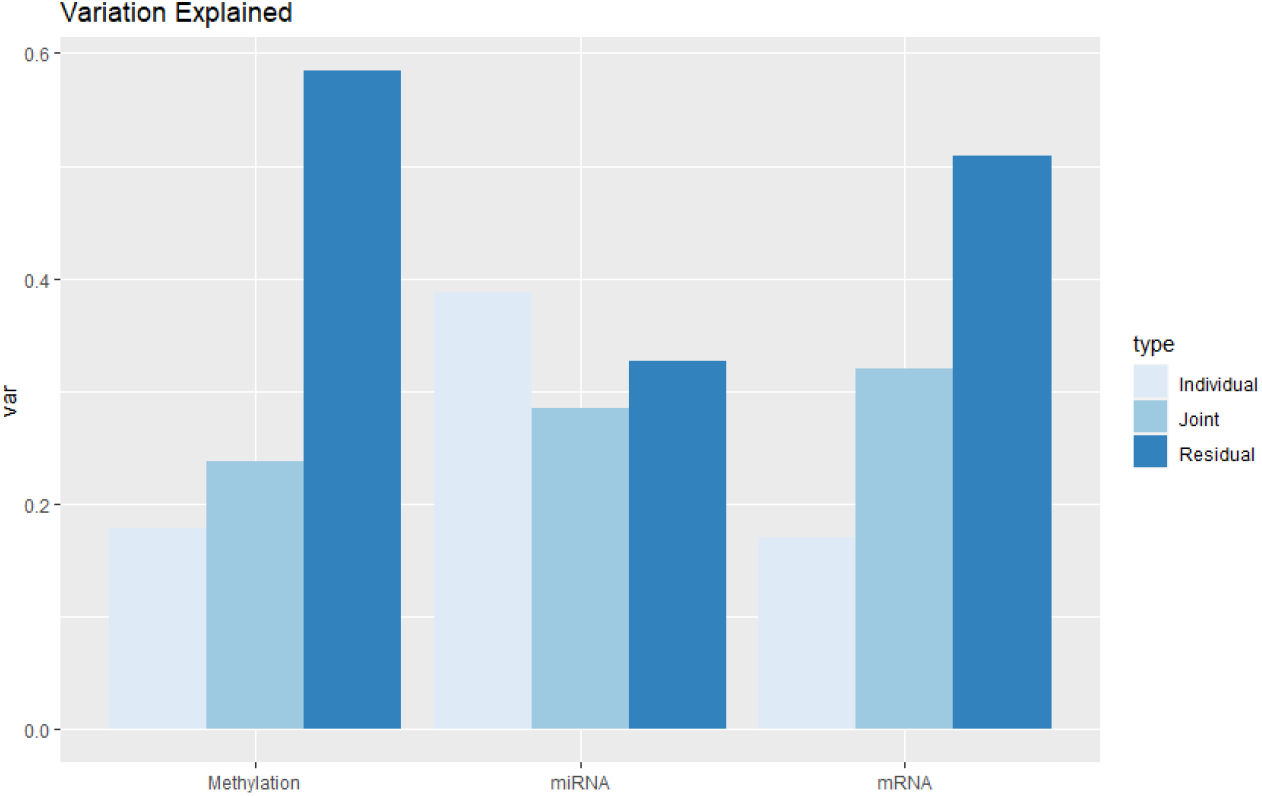
Joint and individual proportions of variance explained in the TCGA dataset. Both joint and individual components are relevant for the three datasets.

#### 3.2.2 Prediction models

An integrative model for prediction of tumor subtype is fitted using the aJIVE joint components and the first five aJIVE individual components for each data source as explanatory variables. This is compared to the non-integrative model, using the first five individual PCs obtained for each data source separately. Multinomial logistic models were used, with four classes for the response variable.

The results were validated by 10-fold cross-validation. Multiclass AUCs for in-sample classification of tumor type, as well as mean AUCs from crossvalidation, are reported in Table 2 for each model. The integrative model including both individual and joint components shows the best quality of prediction.

**Table 2:** In-sample and cross validated AUCs for prediction of tumor subtype in the TCGA dataset.

| Model | In-sample AUC | mean AUC from CV |
| --- | --- | --- |
| Full integrative aJIVE | 0.924 | 0.883 |
| Non-integrative | 0.887 | 0.867 |

In addition, we used random forest of 1, 000 trees to predict tumor subtype on the basis of joint and individual components, and again compared them to non-integrative models. Table 3 reports accuracy and OOB classification error for the random forests, as well as the mean AUCs. The integrative model performs better than the non-integrative model in terms of AUC. In terms of accuracy and classification error, the integrative model and the non-integrative model are equivalent. Figure 5 shows the first ten variables ranked by variable importance in the full integrative model. The three top variables are joint components, and their importance measured in mean gini index is substantially higher than the importance of the other variables.

**Figure 5:**
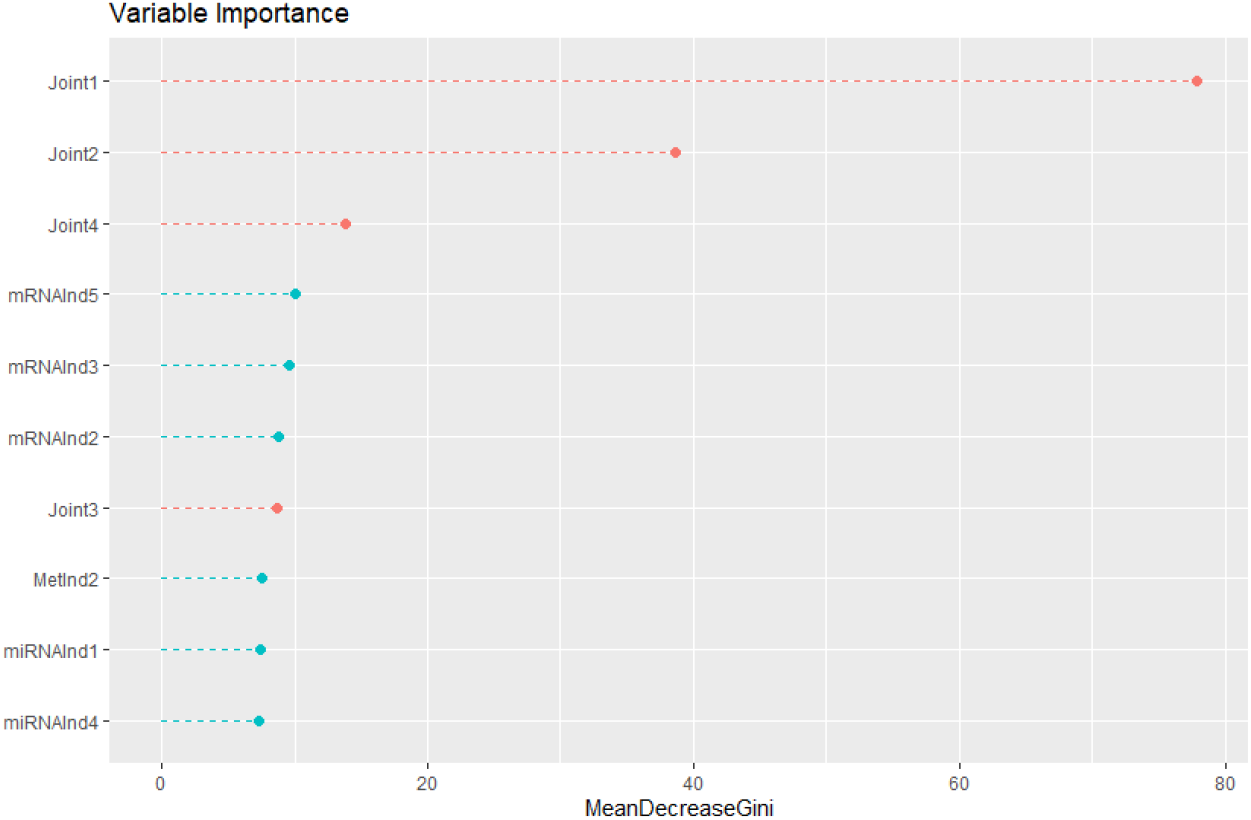
Variable importance plot from random forest on cancer subtype in the TCGA dataset. First ten variables ranked by variable importance (in terms of mean Gini index) in the full integrative model for cancer subtype in the TCGA dataset. `Joint`_*i*_ denotes the *i*—th joint component estimates by aJIVE, while `MetInd`_*i*_, `mRNAInd`_*i*_ and `miRNAInd`_*i*_ are the *i*—th individual components estimated by aJIVE for methylation, mRNA and miRNA respectively.

**Table 3:** Random forest diagnostics for prediction models of cancer subtype in the TCGA dataset.

| Model | Accuracy | OOB Classification Error | AUC |
| --- | --- | --- | --- |
| Full integrative aJIVE | 0.81 | 19% | 0.91 |
| Non-integrative | 0.81 | 19% | 0.78 |

## 4 Discussion

### Prediction results

We use data integration to identify both joint and individual components in a lung cancer study, where multiple omics data sources are available. While the individual contribution of each data source is known to be relevant and has been widely studied in this context, different data sources are also expected to jointly associate with the clinical outcomes. We show how including both joint and individual components in prediction models improves the quality of prediction of the occurrence of lung cancer, as well as its classification into metastatic or non-metastatic cancer. Models that include both types of components lead to better predictions when compared to non-integrative models, or to models based on clinical covariates. This approach was also used on data from a breast cancer study available from The Cancer Genome Atlas (TCGA Research Network) and the results are similar.

Prediction models are validated in a 10-fold cross-validation framework, and such results are further confirmed by random forests. From the crossvalidation study, we see that for case-control status, the integrative analysis provides better prediction than non-integrative analysis.

As an additional comparison, we use supervised variable selection and fit a lasso model on the three omics layers. Although we would expect that a supervised method, by using information from the response variable in fitting the model itself, results in better predictions, the aJIVE models used here perform similarly (for case vs control), and substantially better (for metastasis).

A possible explanation of generally low AUCs is that prediction models for the NOWAC dataset might also be affected by the time between blood sampling and cancer diagnosis, and we expect the quality of predictions to be higher in subjects with a shorter time to diagnosis. We stratified cases into two subgroups based on time to diagnosis (higher vs lower than the median time) and obtained higher in-sample AUCs for the classification of case vs control in subjects with a closer time to diagnosis. For the classification of metastasis, the sample size in the two time to diagnosis classes is not enough to draw conclusions. In the application to the NOWAC dataset, it is interesting to observe that one genomic component identified from aJIVE, specifically one individual component from the methylation data, ranks above smoking in importance for case-control classification (Figure 3), smoking being known to be the one, major risk factor for lung cancer.

In the TCGA example, the high prediction quality of the non-integrative model is likely to arise from the definition of subtypes, which is based on levels of mRNA expression. In the non-integrative models, the mRNA principal components highly contribute to the prediction quality (mean AUC from 10-fold CV = 0.875 for non-integrative models based on mRNA components only vs mean AUC from 10-fold CV = 0.867 for the non-integrative models based on all three sources). Although we use multiclass ROC curves in this example, a dichotomous classification of the tumor classes could provide a deeper understanding of the models and easier comparison with the logistic case.

While this work provides preliminary evidence of the importance of an integrative analysis of the omics sources, a more thorough investigation of the joint and individual components could help identifying relevant biological patterns for future research. An example can be given by the underlying biological processes involving smoking and lung cancer: the omics signals that are dominating the components could be important risk factors for lung cancer, in addition to being informative about present or past smoking, and their interaction could shed light on the relevant underlying biological processes. Although a functional interpretation of such processes and of their link to the clinical outcomes is not straightforward, an investigation of the aJIVE components could provide further information that would not be identified by a non-integrative analysis of the separate omics sources.

### Variable filtering

The chosen approach for variable filtering is based on variance for mRNA, and on genomic location and variance for methylation. Specifically, the top 5,000 most variable mRNAs are selected and CpGs are then selected based on their location on genes, by including CpGs located on the same genes as filtered mRNAs. The top 10,000 most variable CpGs are included in addition to these. We expect that choosing signals that are on the same gene locations, and therefore naturally associated, will result in very relevant joint contributions and possibly obscure the individual components associated with methylation. The inclusion of the most variable CpGs, independently from their gene locations, solves this issue. Varying the proportions of CpGs that are selected based on their variance and on their gene location can give rise to different joint and individual contributions, and this aspect needs to be fully considered in the interpretation of the results. In the supplementary material (File 2), we report the aJIVE results for two additional filtering set ups, specifically: a) by selecting CpGs uniquely on the basis of their gene location, and b) by including only the top most variable CpGs regardless of their location. The filtering of mRNA is based on the variance of the log-transformed signals. Although this procedure might generally result in selecting signals with the lowest intensities, this did not seem to have any impact on the results in our example. Different choices of filtering criteria for the mRNAs can be the interquantile range (IQR), or the association with the clinical outcome of interest, estimated by an appropriate regression model, and would yield different results of the aJIVE decomposition. Finally, also the filtering of the miRNAs needs to be taken into account, where less restrictive criteria might result in the estimation of different joint and individual components. Other choices could be made in this phase, for example applying the variance criterion independently on each data source, which could yield different joint and individual components. Another choice we made in the preprocessing and filtering of the data is the use of M-values for methylation. This choice is motivated by Du et al (2010).

### Methodological considerations

One of the main issues in aJIVE is the selection of initial ranks. The most common method for the choice of initial ranks in aJIVE is the visualization of screeplots, which is subjective and highly sensitive to noise in the data. The profile likelihood idea suggested by Zhu and Ghodsi (2006) partly addresses the problem, but it still lacks some objectivity and automation. Nevertheless, the correct choice of ranks is fundamental for aJIVE, and ranks misspecification can lead to incorrect results (Feng et al, 2018).

The high dimensionality of the data motivates the use of sparse methods, which reduce the number of variables included in the model and provide an easier interpretation of the results. A sparse version of the aJIVE method could be used for this purpose, by introducing a penalty term in the decomposition to induce variable sparsity. This has not been specifically implemented for aJIVE, but Lock et al (2013) discuss and provide an implementation of a sparse version of the JIVE method.

Finally, one aspect that is not accounted for in aJIVE is the presence of partially shared components. When joint components are only shared by, for example, two out of the three data sources, they will not be identified by aJIVE. This is a limitation of most data integration methods, and we expect partially shared components to result in even better prediction models. A way to investigate partially shared patterns is provided in the SLIDE method by Gayananova and Li (2019), and is a potential starting point for further work in this direction.

## 5 Conclusion

Our study shows how integrative models that include both joint and individual contribution of multiple datasets lead to more accurate model predictions, and facilitate the interpretation of the underlying biological processes. We use joint and individual contributions of DNA methylation, miRNA and mRNA expression to predict cancer development in a lung cancer case-control study, and breast cancer subtype in a dataset from The Cancer Genome Atlas. We show that the use of joint and individual components leads to better prediction models, and to a deeper understanding of the biological process in hand.

## Ethics approval and consent to participate

All participants gave written informed consent and the study was approved by the Regional Committee for Medical and Health Research Ethics and the Norwegian Data Inspectorate. More information is available in Lund et al (2008).

## Availability of data and materials

The code for the statistical analysis is available at https://github.com/ericaponzi.

Data cannot be shared publicly because of local and national ethical and security policies. Data access for researchers will be conditional on adherence to both the data access procedures of the Norwegian Women and Cancer Cohort and the UiT – The Arctic University of Norway (contact via Tonje Braaten and Arne Bastian Wiik) in addition to an approval from the local ethical committee.

## Acknowledgements

The miRNA and mRNA analyses were provided by the Genomics Core Facility (GCF), Norwegian University of Science and Technology (NTNU). GCF is funded by the Faculty of Medicine and Health Sciences at NTNU and Central Norway Regional Health Authority.

## Author’s contributions

K. M., M. T. and E. P. conceived the research idea. E.P. conducted the statistical analyses. T. H. N. was responsible for the acquisition of data and the biological interpretation of the results. E.P. K. M. and M.T. wrote the manuscript, with inputs from T. H. N. All authors gave final approval.

