## Supplementary File 1 for "Integrative, multi-omics, analysis of blood samples improves model predictions: applications to cancer"

### Supplementary material for “Integrative, multi-omics, analysis of blood samples improves model predictions: applications to lung and breast cancer”

##### Different filtering methods on the NOWAC dataset

We examine different filtering criteria and report aJIVE results on the NOWAC dataset in the different cases. In the first case, the filtering of CpGs is based on their gene location, and we pick those CpGs that are located on the same genes as the selected mRNAs. This is expected to cause high joint components in the aJIVE estimation. In the second case, we focus on the individual contributions and select CpGs based on their variance and regardless of their gene location.

#### 1 Filtering based on gene location

We reduced the number of mRNA expressions to $p_2 = 5000$, by selecting the variables with higher variance. We then reduced the number of CpGs methylation sites by selecting the CpGs located on the same genes as the filtered mRNAs. Among these, we excluded CpGs with more than 40% missing data, as well as CpGs with extreme M-values (see main text). This resulted in $p_1 = 18545$. All available miRNAs were included.

##### 1.1 aJIVE results

Using initial ranks obtained with the profile likelihood method resulted in a joint rank equal to 5, and individual ranks equal to 43, 6, and 7, respectively for mRNA, miRNA and methylation. Figure 1 reports the proportions of variance explained that are due to the joint, individual and residual components.

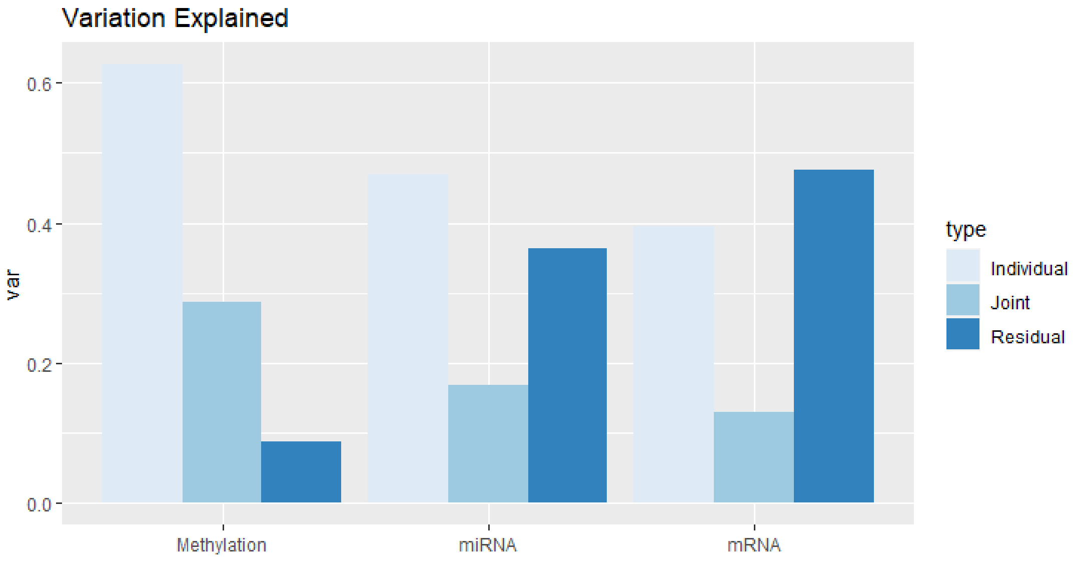


Figure 1: Proportion of variation explained by joint, individual and residual components in each data source for filtering based on gene locations.

#### Prediction models

Figures 2 and 3 report the in-sample ROC curves relative to the logistic models fitted on the joint and individual components estimated by aJIVE. The model with only patient covariates (age, BMI and smoking) as explanatory variables and the full, integrative model are reported. The integrative model is fitted using patient covariates, aJIVE joint components and first five aJIVE individual components for each data source as explanatory variables. These are compared to non-integrative models, using the first five individual PCs obtained for each dataset separately, in addition to the same covariates.

These results were validated by 10-fold cross validation for each outcome. In the ROC studies from cross validation, the model with all components seems to improve the prediction for both case-control and metastasis status. The mean AUCs for the integrative models are 0.71 and 0.69, for case-control and metastasis status respectively. The mean AUC of the non-integrative model, based on the single data PCAs and the clinical covariates, is respectively 0.69 and 0.61, lower than the AUCs obtained in the models using the aJIVE components.

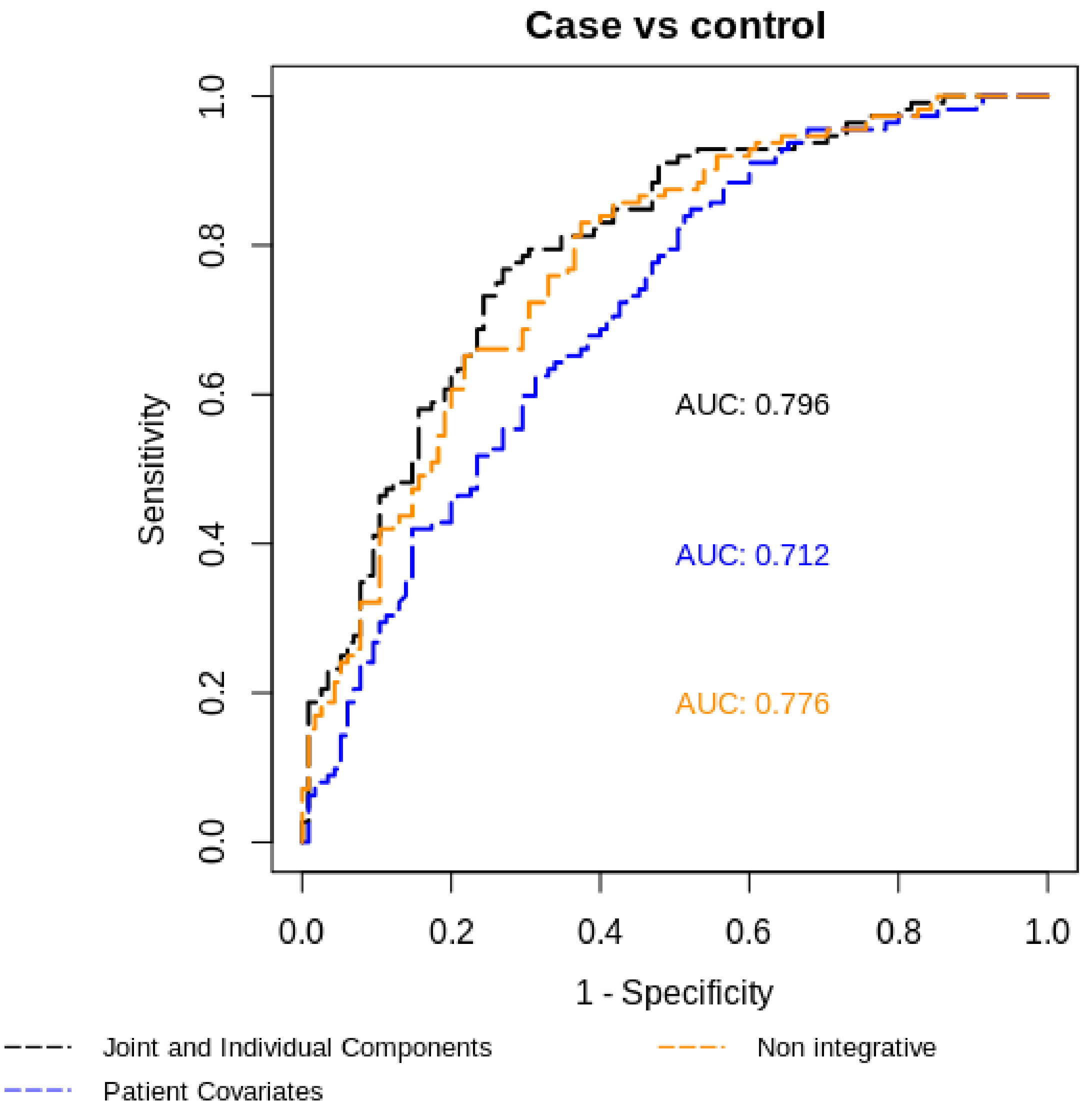


Figure 2: ROC curves for logistic prediction models on case vs control: in-sample predictions

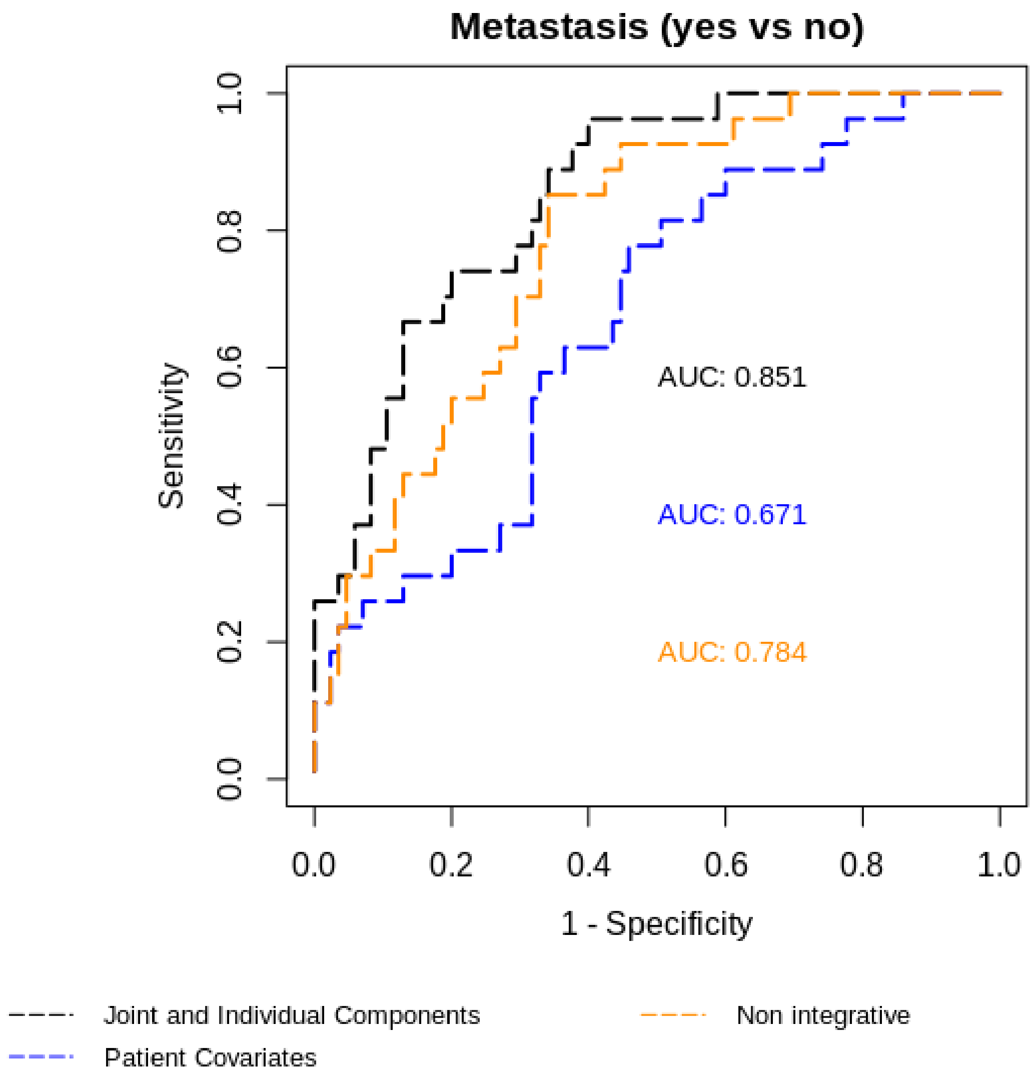


Figure 3: ROC curves for logistic prediction models on metastasis: in-sample predictions

#### 2 Filtering based on variance

We reduced the number of mRNA expressions to $p_2 = 5000$, by selecting the variables with higher variance. We then reduced the number of CpGs methylation sites by selecting the top 50000 CpGs with higher variance. Among these, we excluded CpGs with more than 40% missing data, as well as CpGs with extreme M-values (see main text). This resulted in $p_1 = 46195$. All available miRNAs were included.

#### 2.1 aJIVE results

Using initial ranks obtained with the profile likelihood method resulted in a joint rank equal to 5, and individual ranks equal to 47, 7 and 6, respectively for mRNA, miRNA and methylation. Figure 4 reports the proportions of variance explained that are due to the joint, individual and residual components.

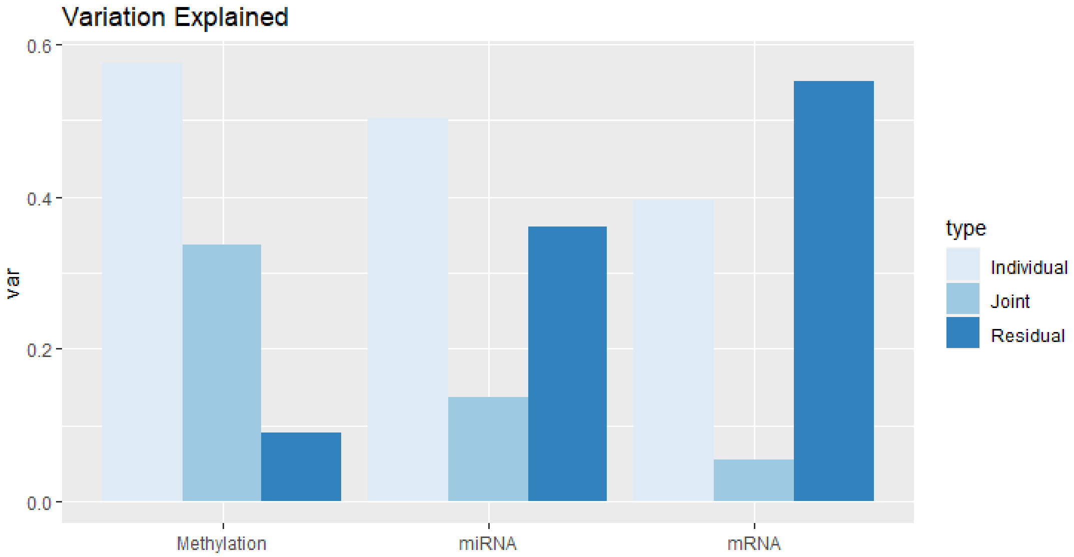


Figure 4: Proportion of variation explained by joint, individual and residual components in each data source for filtering based on variation.

#### Prediction models

Figures 5 and 6 reports the in-sample ROC curves relative to the logistic models reported above.

These results were validated by 10-fold cross validation for each outcome. In the ROC studies from cross validation, the integrative models improve the prediction for both case-control and metastasis status. The mean AUCs for the integrative models are 0.68 and 0.74, for case-control and metastasis status respectively. The mean AUC of the non-integrative model, based on the single data PCAs and the clinical covariates, is respectively 0.65 and 0.69.

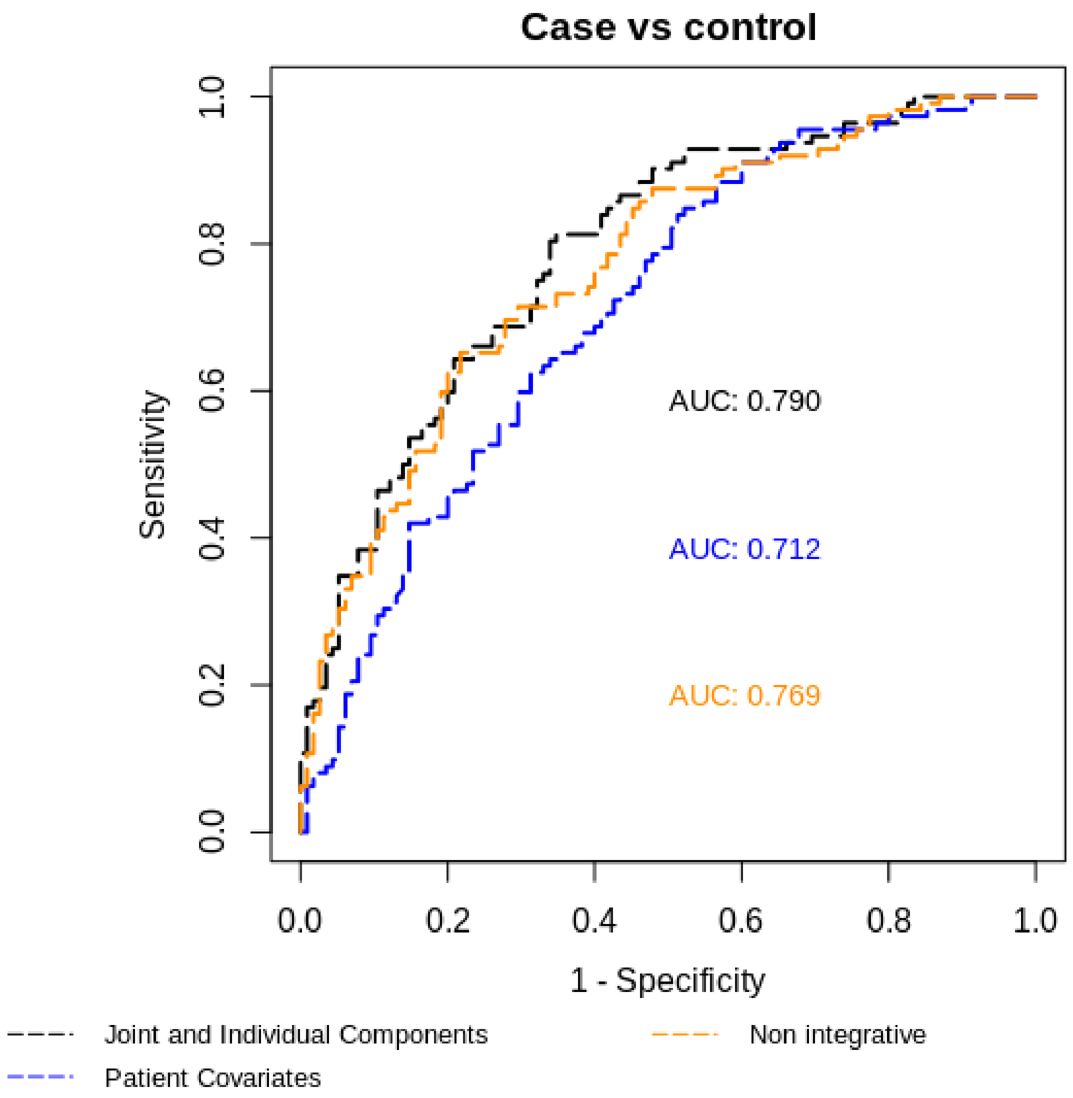


Figure 5: ROC curves for logistic prediction models on case vs control: in-sample predictions

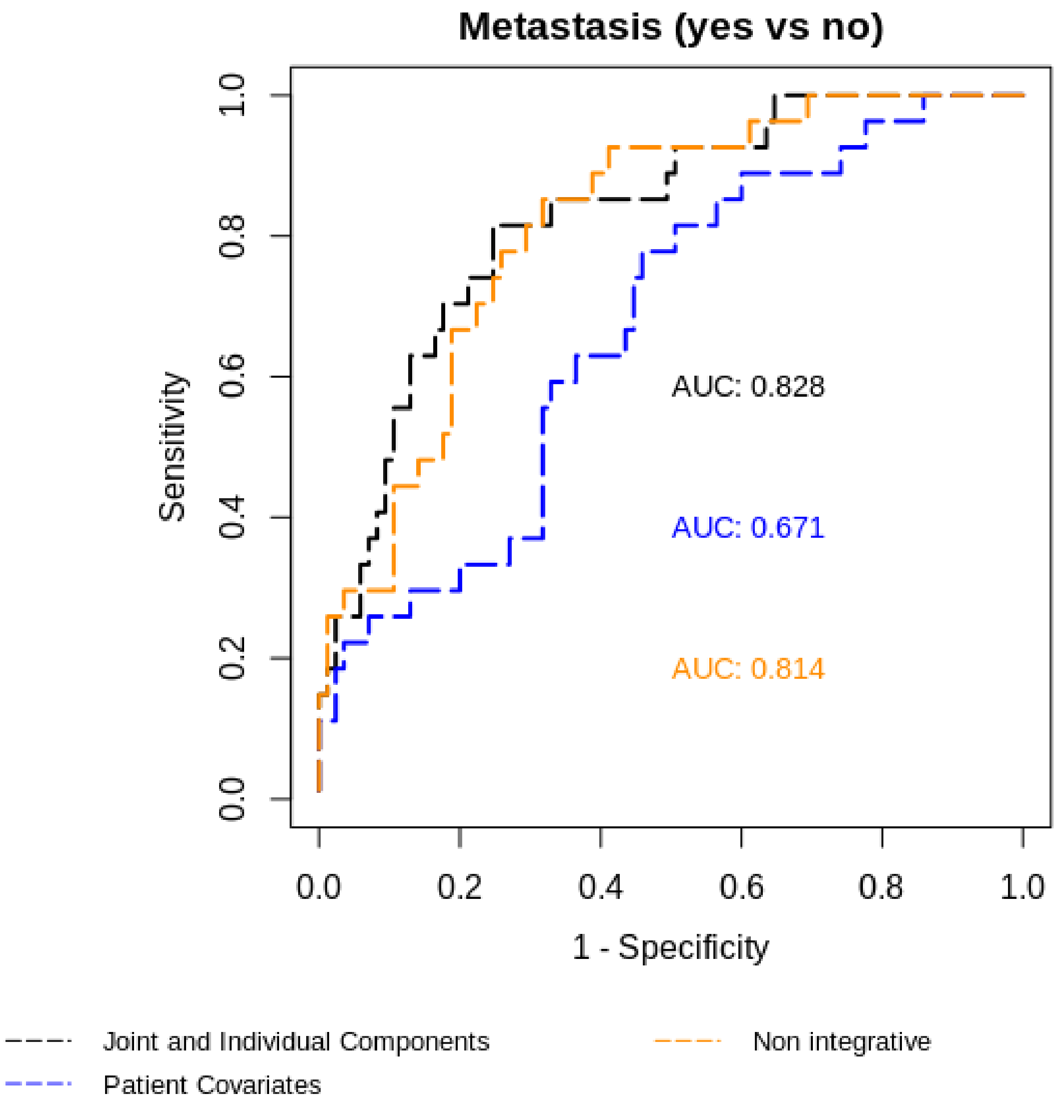


Figure 6: ROC curves for logistic prediction models on metastasis: in-sample predictions
