## Supplementary File 2 for "Integrative, multi-omics, analysis of blood samples improves model predictions: applications to cancer"

### Supplementary material for “Integrative, multi-omics, analysis of blood samples improves model predictions: applications to lung and breast cancer”

#### Effect of initial ranks on the proportion of variance explained

We examine different choices of initial ranks and report the estimated proportion of variance explained for the NOWAC dataset. We set initial ranks respectively lower and higher than the ones estimated by profile likelihood, and report the proportions of joint, individual and residual variance explained for each data source.

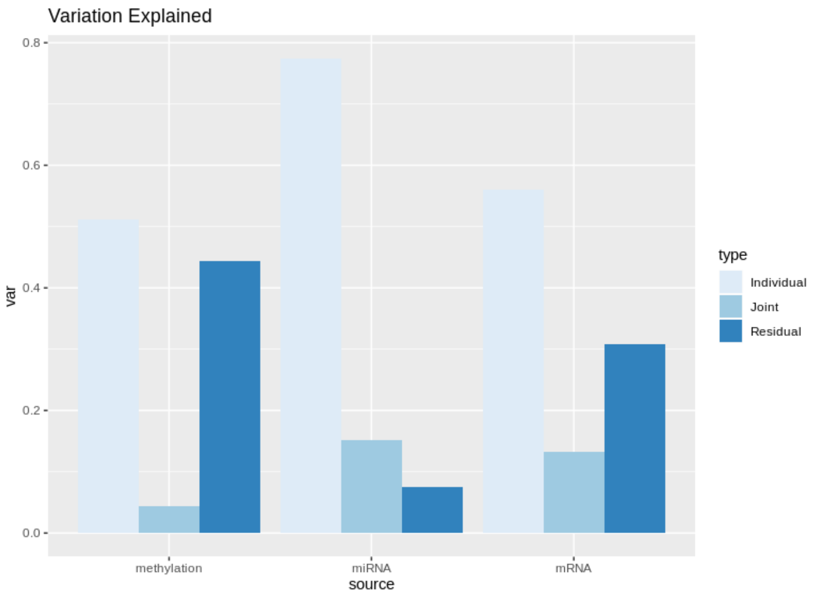


Figure 1: Proportions of variance explained with low initial ranks.

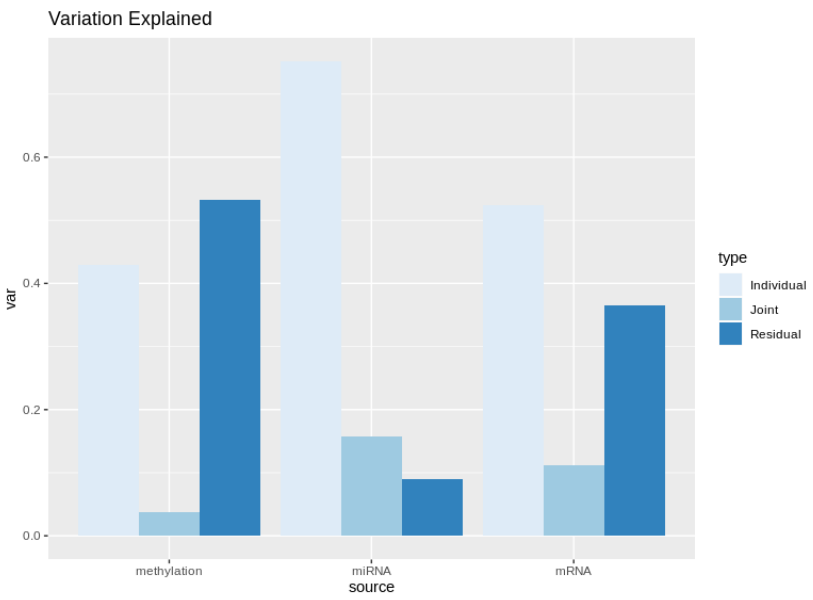


Figure 2: Proportions of variance explained with high initial ranks.

The choice of initial ranks has a marginal effect on the estimated proportions of variance explained. As pointed out in (1), rank misspecification affects the aJIVE estimation when the misspecified ranks are lower than the true ranks, while using higher ranks does not alter the aJIVE results substantially.
